# Genome sequence of the corn leaf aphid (*Rhopalosiphum maidis* Fitch)

**DOI:** 10.1101/438499

**Authors:** Wenbo Chen, Sara Shakir, Mahdiyeh Bigham, Zhangjun Fei, Georg Jander

## Abstract

**Background:** The corn leaf aphid (*Rhopalosiphum maidis* Fitch) is the most economically damaging aphid pest on maize (*Zea mays*), one of the world’s most important grain crops. In addition to causing direct damage due to the removal of photoassimilates, *R. maidis* transmits several destructive maize viruses, including *Maize yellow dwarf virus*, *Barley yellow dwarf virus*, *Sugarcane mosaic virus*, and *Cucumber mosaic virus*.

**Findings:** A 326-Mb genome assembly of BTI-1, a parthenogenetically reproducing *R. maidis* clone, was generated with a combination of PacBio (208-fold coverage) and Illumina sequencing (80-fold coverage), which contains a total of 689 contigs with an N50 size of 9.0 Mb. The contigs were further clustered into four scaffolds using the Phase Genomics Hi-C interaction maps, consistent with the commonly observed 2n = 8 karyotype of *R. maidis*. Most of the assembled contigs (473 spanning 321 Mb) were successfully orientated in the four scaffolds. The *R. maidis* genome assembly captured the full length of 95.8% of the core eukaryotic genes, suggesting that it is highly complete. Repetitive sequences accounted for 21.2% of the assembly, and a total of 17,647 protein-coding genes were predicted in the *R. maidis* genome with integrated evidence from *ab initio* and homology-based gene predictions and transcriptome sequences generated with both PacBio and Illumina. An analysis of likely horizontally transferred genes identified two from bacteria, seven from fungi, two from protozoa, and nine from algae.

**Conclusions:** A high-quality *R. maidis* genome was assembled at the chromosome level. This genome sequence will enable further research related to ecological interactions, virus transmission, pesticide resistance, and other aspects of *R. maidis* biology. It also serves as a valuable resource for comparative investigation of other aphid species.

## Data Description

### Introduction

Maize (*Zea mays*), the world’s most productive grain crop, is susceptible to more than 90 species of herbivorous insects [1-3]. Among aphids that feed on maize, the corn leaf aphid (*Rhopalosiphum maidis* Fitch) is the most commonly encountered, particularly in tropical and warmer temperate areas [4]. Relative to other maize-feeding aphids (*Rhopalosiphum padi*, *Schizaphis graminum*, *Sitobion avenae*, and *Metopolophium dirhodum*), *R. maidis* exhibits a greater tolerance of benzoxazinoids, the most abundant class of maize defensive metabolites [5]. However, the mechanism of aphid resistance to these plant toxins is not known, and natural variation in benzoxazinoid content among maize inbred lines nevertheless influences growth and reproduction of *R. maidis* [6, 7].

Damage caused to maize by *R. maidis* takes several forms, and the resulting yield losses can be quite variable from year to year. Growth and yield are reduced through the removal of photosynthates by large numbers of aphids [8]. On flowering-stage maize, aphids tend to congregate on the tassels, where large amounts of honeydew can prevent the release of pollen from the anthers, thereby reducing seed set by up to 90% [9, 10]. Additional damage comes from the fact that *R. maidis* transmits several important maize viruses, including *Maize yellow dwarf virus*, *Barley yellow dwarf virus, Sugarcane mosaic virus*, and *Cucumber mosaic virus* [11-15].

In addition to feeding on maize, *R. maidis* also infests a variety of other monocot species, including barley, oat, rice, rye, sorghum, sugarcane, and wheat [4]. In one study, barley was reported as the most suitable grain crop host for *R. maidis* [16]. However, as in the case of maize, there is also considerable within-species variation for *R. maidis* resistance in barley [17].

The origin of *R. maidis* is likely in Asia, and it has been subsequently introduced in most grain-growing areas of the world. In almost all parts of its range, *R. maidis* is anholocyclic. However, sexual reproduction has been reported in Pakistan and Korea, with *Prunus* ssp. as the primary host [18, 19]. In populations in Japan and Kenya, males but not sexually reproducing females have been found [20, 21]. Consistent with the sometimes permanently parthenogenetic life cycle of *R. maidis*, there is within-species variation in the chromosome numbers. Karyotypes of 2n = 8, 9, and 10 have been reported. There also is evidence of host specificity among the karyotypes. Whereas *R. maidis* strains on maize tend to have 2n = 8, those on barley generally have 2n = 10 [22, 23].

Here we report the genome sequence of *R. maidis* isolate BTI-1. Comparisons to six previously published aphid genomes [24-31] showed an improved assembly, with most of the sequences assembled into four scaffolds, consistent with the 2n = 8 karyotype of *R. maidis*. Analysis of the assembled *R. maidis* genome identified horizontally transferred genes, repetitive elements, and likely xenobiotic detoxification enzymes.

### Sampling and genome sequencing

#### Insect Colony

BTI-1, a corn leaf aphid (*R. maidis*) isolate, which was originally collected from maize (*Z. mays*) in New York State, was obtained from Stewart Gray (USDA Plant Soil and Nutrition Laboratory, Ithaca, NY). An isogenic colony was started from a single parthenogenetic female *R. maidis* and was maintained on barley (*Hordeum vulgare*) prior to the collection of insects for genome and transcriptome sequencing.

Genomic DNA was prepared from 100-200 mg of fresh *R. maidis* tissue using a previously described protocol [32]. Briefly, mixed-instar whole aphids were ground in liquid nitrogen and incubated at 65°C in microprep buffer made up of DNA extraction buffer (0.35M sorbitol, 0.1M Tris-base, pH7.5, 5mM ethylenediaminetetraacetic acid), nuclei lysis buffer (0.2M Tris-base, pH 7.5, 0.05M ethylenediaminetetraacetic acid, 2M NaCl, 2% cetyl trimethylammonium bromide), 5% sarkosyl and 0.5% sodium bisulfite for 30 min. This solution was then treated with chloroform:isoamyl alcohol (24:1) and centrifuged for 10 min at 14,000 *g*. The supernatant was treated with RNase A and DNA was pelleted by centrifugation at 4°C at 14,000 *g* for 10 min. DNA pellet was washed with 100% isopropanol and then with 70% ethanol and dissolved in 50 μl of nuclease free water. Around 50 μg of high molecular weight DNA was prepared for PacBio library construction and sequencing using SMRT Cell template preparation kits (Pacific Biosciences), and sequencing was conducted at the Icahn Institute and Department of Genetics and Genomic Sciences, Icahn School of Medicine at Mount Sinai. A total of 16 SMRT Cells were run on the PacBio Sequel platform, yielding 70 Gb raw sequence data (**Supplemental Table S1**), representing a 208-fold coverage of the *R. maidis* genome, which was estimated to be 338 Mb using the kmer approach ([33]; **Figure 1**). For short-read sequencing, one paired-end library was constructed using the Illumina TruSeq DNA sample preparation kit following the manufacturer’s instructions, and sequenced on an Illumina HiSeq 2500 system, which yielded about 75 Gb of raw sequence data (**Supplemental Table S2**). Raw Illumina reads were processed to remove duplicated read pairs, which were defined as having identical bases in the first 100 bp of both left and right reads, and only one read pair from each duplicated sequence was kept. Illumina adapters and low-quality sequences were removed from the reads using Trimmomatic [34]. The kmer depth distribution of the cleaned high-quality sequences displayed a single peak (**Supplemental Figure S1**), indicating that the sequenced sample has a low level of heterozygosity.

**Figure 1.**
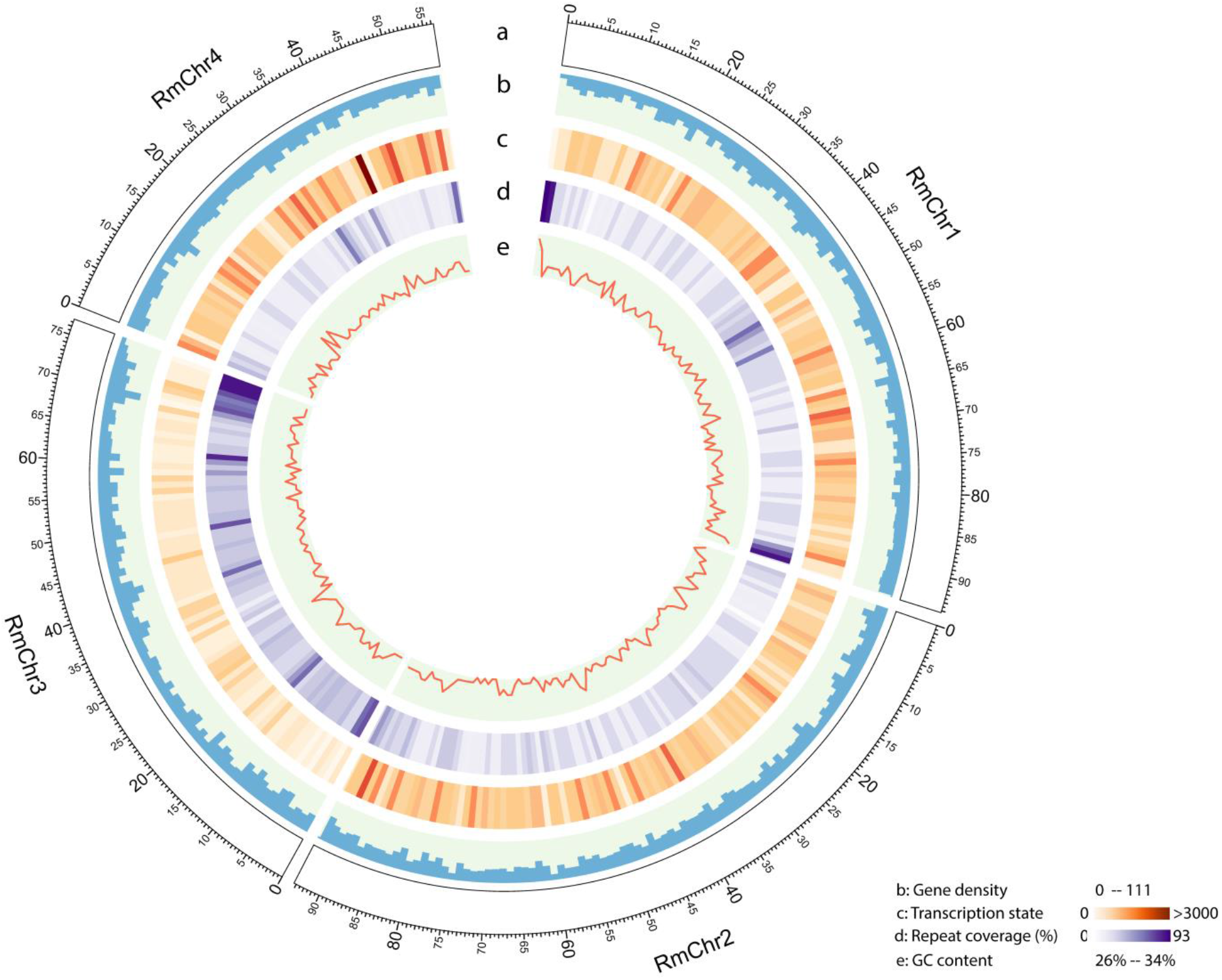
*Rhopalosiphum maidis* genome landscape. (a) Ideogram of the four *R. maidis* pseudochromosomes at the Mb scale. (b) Gene density represented as number of genes per Mb. (c) Transcription state. The transcription level was estimated by read counts per million mapped reads in 1-Mb windows. (d) Percentage of coverage of repeat sequences per Mb. (e) GC content in 1-Mb windows. The four *R. maidis* pseudo-chromosomes represented 98.4% of the genome assembly. This figure was generated using Circos (http://circos.ca/).

### Transcriptome sequencing

Transcriptome sequencing (Illumina strand-specific RNA-Seq and PacBio Iso-Seq) was conducted to aid gene prediction. Total RNA was extracted using the SV Total RNA isolation kit (Promega: Catalog number: Z3100). Briefly, cells were lysed by grinding 100-120 mg of insect tissue in liquid nitrogen, followed by incubation at 70°C in RNA lysis buffer (4M guanidine thiocyanate (GTC), 0.01M Tris, pH 7.5, 0.97% β-mercaptoethanol) for 3 min. This solution was then centrifuged for 10 min at 14,000 *g* and the supernatant was passed through a spin column provided with the kit, followed by DNase treatment. RNA was washed with RNA wash solution (60 mM potassium acetate, 10 mM Tris-HCl (pH 7.5), 60% ethanol) and dissolved in 50 μl of nuclease-free water. Strand-specific RNA-Seq libraries were constructed using a previously described protocol [35] and sequenced at Biotechnology Resource Center of Cornell University on an Illumina HiSeq 2500 sequencing system. More than 188 million paired-end reads with lengths of 151 bp were obtained (**Supplemental Table S2**). Raw reads were processed by trimming adaptor and low-quality sequences using Trimmomatic [34]. The cleaned reads were aligned to the assembled *R. maidis* genome using HISAT2 [36], followed by reference-guided assembly using StringTie [37]. The assembled transcripts were used to improve protein-coding gene predictions in the *R. maidis* genome.

For Iso-Seq, 20 μg RNA, isolated from 100-120 mg of fresh *R. maidis* tissue using the SV Total RNA isolation kit (Promega) with the method described above, was shipped to Duke Center for Genomic and Computational Biology for PacBio large-insert (15-20kb) library construction and sequencing using standard SMRTbell template preparation kits. One SMRT cell was run on the PacBio Sequel platform, yielding ~10 Gb raw sequence data (**Supplemental Table S1**). The PacBio raw reads were processed using IsoSeq3 (https://github.com/PacificBiosciences/IsoSeq3). Briefly, one representative Circular Consensus Sequence (CCS) was generated for each zero-mode waveguide (ZMW). Only ZMWs with at least one full pass, meaning that each primer has been seen at least once, were used for the subsequent analysis. The CCSs were processed to remove the 5’ and 3’ primers, trim off polyA tails and remove artificial concatemers to create full-length, non-concatemer (FLNC) reads. The FLNC reads were then clustered together. The final polishing step created a consensus sequence for each clustered transcript. A total of 21,114 high quality transcripts were generated, and were used to support protein-coding gene predictions in the *R. maidis* genome.

### Hi-C library construction and sequencing

For Hi-C sequencing, 200 mg of *R. maidis* tissue was used for chromatin isolation and library preparation using the animal Hi-C kit from Phase Genomics (https://phasegenomics.com). Hi-C Libraries were sequenced at the Biotechnology Resource Center, Cornell University, using the NextSeq500 platform (Illumina) to obtain 76-nt paired-end reads. Raw reads were processed by trimming adaptor and low-quality sequences using Trimmomatic [34]. The cleaned Hi-C reads were aligned to the assembled contigs using BWA-aln [38], and the optimal placement of each read pair was determined by BWA-sampe [38]. Reads that did not map within 500 bp of a restriction enzyme site were removed using the PreprocessSAMs.pl script in LACHESIS [39]. Finally, only reads with mapping quality greater than 30 were used for scaffolding by LACHESIS [39].

### Genome assembly

The PacBio long reads were corrected and assembled with the Canu assembler [40] (version 1.6). The resulting contigs were polished by aligning the raw PacBio reads to the assembly, and correcting the sequencing errors using Arrow (https://github.com/PacificBiosciences/GenomicConsensus). To further improve the assembly, another round of polishing was performed by aligning the Illumina short reads to the assembly and correcting the sequencing errors using Pilon [41]. The assembled contigs were then compared against the NCBI non-redundant nucleotide (nt) database using BLASTN with an e-value cutoff of 1e-5. Contigs with over 90% of their lengths similar to only bacterial or viral sequences were considered to be contaminants and were discarded. The final contigs were clustered and ordered into chromosomes by Hi-C reads using LACHESIS [39] with default parameters. Scaffolds were manually polished using Juicebox [42].

The assembled *R. maidis* genome had a total length of 326.0 Mb and consisted of 689 contigs with an N50 length of 9.0 Mb. Thus, this is a much-improved genome assembly compared to the six previously published aphid genomes (**Table 1**). A total of 602 contigs spanning 323.4 Mb (99.2% of the assembly) were clustered into four groups, which was consistent with the commonly observed 2n = 8 karyotype of *R. maidis* [22]. Of the clustered contigs, 473 spanning 320.6 Mb (98.4% of the assembly) were successfully orientated (**Figure 1, Supplemental Figure 2**). To evaluate the completeness of the *R. maidis* genome assembly, the Illumina paired-end library were aligned to the assembly, allowing up to three mismatches using BWA-MEM [38]. With this approach, 94.9% of the Illumina reads could be mapped back to the assembly, indicating that most of the reads were successfully assembled into the genome. RNA-Seq reads also were aligned to the genome assembly using HISAT2 [36], resulting a mapping ratio of 94.5% (**Supplemental Table S2**). Furthermore, the completeness of the genome assembly, as evaluated by BUSCO (v 3.0.2 [43], showed that 95.8% of the core eukaryotic genes were at least partially captured by the genome assembly and 94.5% were completely captured. Taken together, our evaluations indicated an overall high quality of the assembled *R. maidis* genome.

**Table 1.** Assembly statistics of seven aphid genomes.

|  | *R. maidis* | *Ap. glycines* | *M. persicae* | *Ac. pisum* | *M. cerasi* | *R. padi* | *D. noxia* |
| --- | --- | --- | --- | --- | --- | --- | --- |
| Genome assembly |  |  |  |  |  |  |  |
| Assembly size (Mb) | 326.0 | 302.9 | 347.3 | 541.6 | 405.7 | 319.4 | 393.0 |
| Contig Count | 689 | 66,000 | 8,249 | 60,623 | 56,508 | 16,689 | 49,357 |
| Contig N50 (bp) | 9,046,396 | 15,844 | 144,275 | 28,192 | 17,908 | 96,831 | 12,578 |
| Scaffold Count | 220 | 8,397 | 4,022 | 23,924 | 49,286 | 15,587 | 5,641 |
| Scaffold N50 (bp) | 93,298,903 | 174,505 | 435,781 | 518,546 | 23,273 | 116,185 | 397,774 |
| Max. scaffold length (Mb) | 94.2 | 1.4 | 2.2 | 3 | 0.2 | 0.6 | 2.1 |
| Min. scaffold length (kb) | 1.1 | 2 | 0.9 | 0.2 | 1 | 1 | 0.9 |
| Genomic features |  |  |  |  |  |  |  |
| Gene count | 17,647 | 19,182 | 18,529 | 36,195 | 28,688 | 26,286 | 25,987 |
| Transcript length (bp) | 1,834.6 | 1,520.1 | 1,838.7 | 1,964.1 | NA | NA | NA |
| CDS length (bp) | 1,242.04 | 1,240.3 | 1,328.3 | 1,157.6 | 952.7 | 1155.09 | 970.2 |
| exon length (bp) | 210.02 | 245.5 | 299.2 | 394.7 | NA | NA | NA |
| exon count/gene | 6.31 | 6.19 | 6.14 | 4.97 | NA | NA | NA |

NA: This information could not be retrieved from the annotation files.

### Endosymbiont genomes

The genome sequence of the *Buchnera aphidicola* endosymbiont was separated from the *R. maidis* host genome sequences by aligning the initial assembly to the *Buchnera* reference genome (GeneBank ID: NC_002528.1). One single contig was extracted and polished using both PacBio long reads and Illumina short reads, as described above. Genome annotation was performed using prokka [44]. The assembled *Buchnera*Rm genome had a length of 642,929 bp (**Supplemental Figure S3**), with a total of 602 predicted protein-coding genes. The two *Buchnera* plasmids, pLeu and pTrp, were also sequenced and assembled, with lengths of 7,852 bp and 3,674 bp, respectively (**Supplemental Figure S3**).

To identify secondary bacterial symbionts in *R. maidis*, raw assembled contigs were compared against the reference sequences of previously identified secondary bacterial symbionts of aphids, including *Hamiltonella defensa*, *Regiella insecticola*, *Serratia symbiotica*, *Rickettsia*, *Spiroplasma*, X-type, *Arsenophonus*, and *Wolbachia* [45], using BLAST. No hits were found, suggesting that these secondary bacterial symbionts are not hosted by the sequenced *R. maidis* strain.

### Annotation of repetitive elements

We first identified MITE (miniature inverted-repeat transposable elements) from the assembled *R. maidis* genome using MITE-Hunter [46], and then generated a *de novo* repeat library by scanning the assembled genome using RepeatModeler (http://www.repeatmasker.org/RepeatModeler), which integrates results from RECON [47], TRF [48], and RepeatScout [49] and classifies repeats with the RepBase library [50]. RepeatModeler identified a total of 546 repeats. We subsequently compared these repeat sequences against the NCBI non-redundant (nr) protein database using BLAST with an e-value cutoff of 1e-5, and those having hits to known protein sequences were excluded. Finally, we identified repeat sequences by scanning the assembled *R. maidis* genome using the *de novo* repeat library with RepeatMasker (http://www.repeatmasker.org/) and the RepeatRunner subroutine (http://www.yandell-lab.org/software/repeatrunner.html) in the MAKER annotation pipeline [51]. A total of 21.2% of the assembled *R. maidis* genome was annotated as repeat elements (**Table 2**). The most predominant repeat elements were unknown repeats and MITEs, which occupied 5.6% and 4.4% of the genome respectively.

**Table 2.** Repeats in the *R. maidis* genome assembly

| Class | No. of Copies | Length (bp) | Coverage of genome (%) |
| --- | --- | --- | --- |
| SINE | 27,308 | 7,085,803 | 2.17 |
| LINE | 6,688 | 1,596,259 | 0.49 |
| LTR | 3,445 | 896,470 | 0.28 |
| DNA transposon | 53,797 | 9,499,710 | 2.91 |
| MITE | 64,663 | 14,240,430 | 4.37 |
| Unclassified | 49,627 | 18,375,079 | 5.64 |
| Other* | 375,149 | 17,360,944 | 5.33 |
| Total | 580,677 | 69,054,695 | 21.18 |

\*Other: microsatellites/simple repeats/low complexity sequences

### Gene prediction

Protein-coding genes were predicted from the genome assembly of *R. maidis* using the automated pipeline MAKER [51]. MAKER integrates the results from *ab initio* gene predictions with experimental gene evidence to produce final consensus gene set. The evidence that was used included complete aphid coding sequences collected from NCBI, transcripts assembled from our strand-specific RNA-Seq data, high quality transcript sequences from Iso-Seq, completed proteomes of *Acyrthosiphon pisum, Aphis glycines, Diuraphis noxia, Myzus cerasi, Myzus persicae*, and *Rhopalosiphum padi*, and proteins from the Swiss-Prot database. All of these sequences were aligned to the *R. maidis* genome using Spaln [52]. MAKER was used to run a battery of trained gene predictors, including Augustus [53], BRAKER [54] and GeneMark-ET [55], and then integrated the experimental gene evidence to produce evidence-based predictions. Altogether, 17,647 protein-coding genes were predicted in the *R. maidis* genome. The gene count of *R. maidis* is close to those in *Ap. Glycines* and *M. persicae*, while fewer than those in *Ac. Pisum*, *M. cerasi*, *R. padi* and *D. noxia*, which possess larger genome sizes (**Table 1**). The mean lengths of coding sequences were similar, with the exception of *M. cerasi* and *D. noxia*.

To functionally annotate the predicted genes, their protein sequences were compared against different protein databases including UnitProt (TrEMBL and SwissProt) and two insect proteomes (pea aphid and psyllid) using BLAST with an e-value cutoff of 1e-4. The protein sequences were also compared against the InterPro domain database [56]. GO annotation was performed with Blast2GO [57]. Among the 17,647 predicted *R. maidis* genes, 75.6% had hits to proteins in the Swiss-Prot or TrEMBL database, 36.0% were annotated with GO terms, 75.2% contained InterPro domains, 76.3% shared detectable homology with *A. pisum* genes, and 47.9% shared detectable homology with *Diaphorina citri* genes.

### Comparative genomics

We compared the *R. maidis* genes with those of six other aphid species (*Ap. glycines, M. persicae*, *Ac. pisum, M. cerasi, R. padi*, and *D. noxia*), as well as the whitefly (*Bemisia tabaci*) [24-31]. The proteome sequences of all eight species were used to construct orthologous groups using OrthoMCL [58]. A total of 5,696 orthologous groups were shared by all 16 species, including 3,605 single-copy orthologous genes. These single-copy genes were used to reconstruct their phylogenetic relationships. Briefly, protein sequences of the single-copy genes were aligned with MUSCLE [59], and positions in the alignment containing gaps in more than 20% of the sequences were removed by trimAl [60]. A phylogenetic tree was then constructed using the Maximum-Likelihood method implemented in PhyML [61], with the JTT model for amino acid substitutions and the aLRT method for branch support. *B. tabaci* was used as the outgroup in the phylogenetic tree, which showed that *R. maidis* is close to *R. padi*, and separated from *A. pisum* and *M. persicae* (**Figure 2**), consistent with a phylogeny that was derived using mtCOI [62].

**Figure 2.**
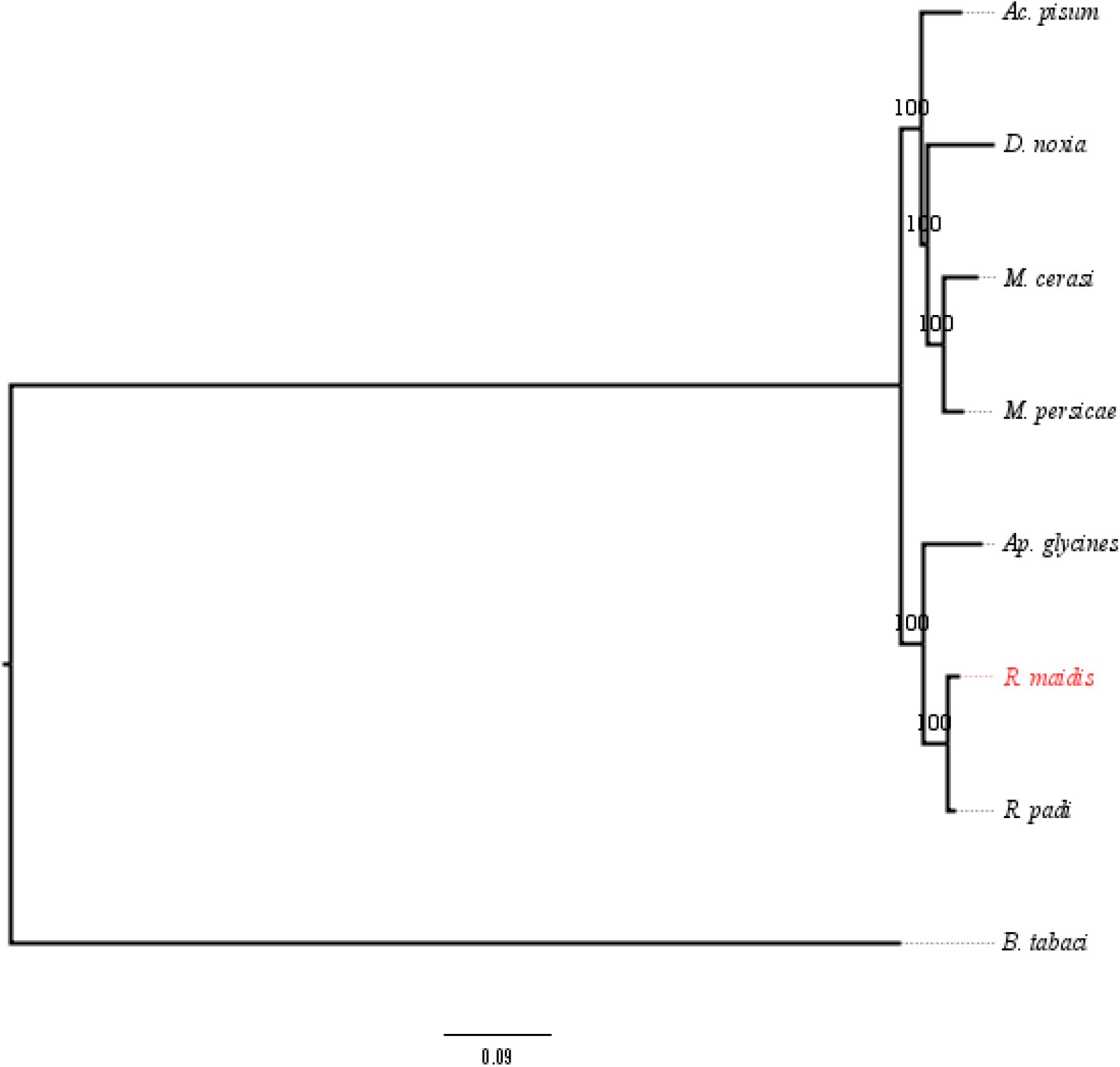
Phylogenetic relationships of *R. maidis* and 7 other arthropod species. *B. tabaci* was used as the outgroup taxon.

### Identification of horizontal gene transfers

All of the *R. maidis* predicted gene models were compared against six protein databases derived from complete proteomes in UniProt, including those from bacteria, archaea, fungi, plants, metazoa (excluding proteins from other species in the Arthropoda), and other eukaryotes, using BLASTP. The index of horizontal gene transfer (HGT), h, was calculated by subtracting the bitscore of the best metazoan match from that of the best non-metazoan match [63]. We required that these sequences were aligned better to the other five taxa than to the metazoan database, defining HGT candidates as those with h≥30 and a best non-metazoan hit bitscore ≥100. The corresponding genome sequences of these candidates as well as 1000-bp flanking sequences at both ends were manually checked for potential genome assembly errors, and none were found.

We phylogenetically validated all HGT candidates. Their protein sequences were compared against the protein databases of six taxa (archaea, bacteria, fungi, plants, metazoan, and other eukaryotes) using BLASTP. The top five hits from each taxon were extracted, and aligned with the candidate HGT protein using ClustalW2 [64]. Each alignment was trimmed to exclude regions where gaps were more than 20% of sequences. Phylogenetic trees were constructed using PhyML [61] using a JTT model with 100 bootstraps. A horizontally transferred gene was considered valid if the gene was monophyletic within the bacteria, archaea, fungi, plants, or protozoa. This analysis identified 20 HGTs, including two of bacterial origin, seven of fungal origin, two from protozoa, and nine from algae (**Table 3**). The two bacterial genes were previously identified as horizontally transferred into *Ac. pisum* [65], and expression silencing of one of these genes, a bacteriocyte-expressed LD-carboxypeptidase A, was shown to reduce aphid performance [66]. A cluster of genes encoding multiple enzymes for carotenoid biosynthesis, which were horizontally transferred into the *Ac. pisum* genome from fungi [67], is also present in the *R. maidis* genome. Two *R. maidis* genes that cluster together with genes from trypanosomes and other protozoa have not been previously reported as horizontally transferred in aphids. Finally, nine genes encoding proteins containing ankyrin repeat domains show highest similarity to genes from unicellular algae in the genus *Ostreococcus*.

**Table 3.** Horizontally transferred genes in *R. maidis*

| Gene ID | Function description | Origin |
| --- | --- | --- |
| Rma07998 | Peptidase U61; LD-carboxypeptidase A | Bacteria |
| Rma09603 | Carbamoylphosphate synthase large subunit | Bacteria |
| Rma01752 | Lycopene cyclase phytoene synthase | Fungi |
| Rma01753 | Carotenoid desaturase | Fungi |
| Rma01754 | Lycopene cyclase phytoene synthase | Fungi |
| Rma01756 | Lycopene cyclase phytoene synthase | Fungi |
| Rma01758 | Lycopene cyclase phytoene synthase | Fungi |
| Rma01759 | Lycopene cyclase phytoene synthase | Fungi |
| Rma01760 | Carotenoid desaturase | Fungi |
| Rma08772 | Leucine Rich Repeat family protein | Protozoa |
| Rma11572 | Antigenic protein, putative | Protozoa |
| Rma10344 | Ankyrin repeat protein | Algae |
| Rma11418 | Ankyrin repeat protein | Algae |
| Rma12243 | Ankyrin repeat protein | Algae |
| Rma13322 | Ankyrin repeat protein | Algae |
| Rma13584 | Ankyrin repeat protein | Algae |
| Rma14036 | Ankyrin repeat protein | Algae |
| Rma15269 | Ankyrin repeat protein | Algae |
| Rma16213 | Ankyrin repeat protein | Algae |
| Rma16838 | Ankyrin repeat protein | Algae |

### Detoxification and insecticide resistance

Cytochrome P450s, glutathione *S-*transferases (GSTs), carboxylesterases, UDP-glucosyltransferases (UGTs), and ABC transporters function in the avoidance and/or detoxification of plant defensive metabolites [68, 69], and insecticide resistance [70, 71]. We identified such detoxification-related genes in *R. maidis* based on protein domains that were predicted through InterProScan [72]. Cytochrome P450 genes were identified if their protein sequences contained the cytochrome P450 domain (InterPro ID: IPR001128). Genes with protein sequences containing the GST N-terminal and/or C-terminal domains (InterPro ID: IPR004045, IPR004046) were identified as GSTs. Carboxylesterases were identified on the basis of protein sequences that contained the carboxylesterase domain (InterPro domain ID: IPR002018) [73]. UDP-glucuronosyltransferases were identified if their protein sequences contained a UDP-glucuronosyl/UDP-glucosyltransferase domain (InterPro domain ID: IPR002213). ABC transporters were identified from the genome if their protein sequences contained an ABC transporter-like domain (InterPro ID: IPR003439). Using the same approach, genes from these families were also identified in the other six aphid genomes (*Ap. glycines*, *M. persicae*, *Ac. pisum*, *M. cerasi*, *R. padi*, *D. noxia*). The number of predicted detoxification genes in *R. maidis* is the lowest among the seven species that were examined (**Table 4** and **Supplemental Table S3**), consistent with *R. maidis* being a specialist monocot herbivore that may require a smaller repertoire of detoxification enzymes. Although the detoxification gene count in *Ac. pisum* was high, the average lengths of the protein sequences were shorter than those in *R. maidis*, *Ap. glycines*, and *M. persicae* (**Supplemental Figure S4**), suggesting that these genes could be incomplete or pseudogenes in *Ac. Pisum*, possibly due to a lower-quality genome assembly.

**Table 4.** Numbers of predicted detoxification genes in seven aphid species

|  | *R.*<br>*maidis* | *Ap.*<br>*glycines* | *M.*<br>*persicae* | *Ac.*<br>*pisum* | *M.*<br>*cerasi* | *R.*<br>*padi* | *D.*<br>*noxia* |
| --- | --- | --- | --- | --- | --- | --- | --- |
| Cytochrome P450s | 59 | 61 | 67 | 82 | 74 | 67 | 60 |
| Glutathione S-transferases | 10 | 12 | 13 | 36 | 12 | 11 | 11 |
| Carboxylesterases | 23 | 31 | 37 | 48 | 36 | 34 | 32 |
| UDP-glucuronosyltransferases | 43 | 47 | 57 | 72 | 48 | 55 | 43 |
| ABC transporters | 68 | 74 | 67 | 126 | 68 | 71 | 63 |
| Total | 203 | 225 | 241 | 364 | 238 | 238 | 209 |

## Conclusion

As the currently most complete aphid genome, our *R. maidis* assembly will provide a valuable resource for comparisons with other species and the investigation of aphid genome evolution. Research on the ecological interactions of *R. maidis*, including host plant choices, detoxification of secondary metabolites, and gene expression responses, will be facilitated by the *R, maidis* genome sequence. Practical applications in agriculture may include the identification of virus transmission mechanisms and new targets for chemical pest control.

## Availability of supporting data

This Whole Genome Shotgun project has been deposited at DDBJ/ENA/GenBank under the accession QORX00000000. The version described in this paper is version QORX01000000. The *Buchnera aphidicola* Rm genome has been deposited in GenBank under accession CP032759. Raw genome and RNA-Seq sequences have been deposited in the NCBI Short Sequence Archive (SRA) under accession SRP164762.

## Supporting information

## Additional Files

**Figure S1**. Kmer (K=31) distribution of Illumina genome sequencing reads of *R. maidis*. The total count of kmers was 11,495,021,417, and the peak of kmer depth was 34. The genome size of *R. maidis* was calculated by dividing the total kmer count by the peak depth, which was approximately 338 Mb.

**Figure S2** Hi-C contact map of the *R. maidis* genome

**Figure S3**. Circular view of the genome of the *Rhopalosiphum maidis* endosymbiont, *Buchnera aphidicola* (A) and its plasmids pLeu (B) and pTrp (C).

**Figure S4**. Length distribution of protein sequences of detoxification gene families in seven aphid species.

**Table S1**. Summary of PacBio long reads

**Table S2**. Summary of Illumina short reads

**Table S3**. Detoxification genes in *Rhopalosiphum maidis*

**Abbreviations**
CCS: circular consensus sequence
ZMW: zero-mode waveguide
FLNC: full-length, non-concatemer reads
nt: nucleotide
HGT: horizontal gene transfer
GST: glutathione S-transferase
CCE: carboxylesterase
UGT: UDP-glucuronosyltransferases

## Author contributions

GJ and ZF conceived of the research, SS and MB raised aphids and isolated nucleic acids, WC conducted data analysis, and WC, SS, and GJ wrote the manuscript.

## Ethics approval and consent to participate

This is not required for experiments with *R. maidis*.

## Acknowledgements

We thank Maximilian Press at Phase Genomics for assistance with the manual polishing of scaffolds using Juicebox.

## Competing interests

The authors have no competing financial and non-financial competing interests.

## Funding

This research was sponsored by the Defense Advanced Research Projects Agency (DARPA) and was accomplished under cooperative agreement number HR0011-17-2-0053. The views and conclusions contained in this document are those of the authors and should not be interpreted as representing the official policies, either expressed or implied, of DARPA or the U.S. Government. The U.S. Government is authorized to reproduce and distribute reprints for Government purposes notwithstanding any copyright notation hereon.

---

**Figure S1.**
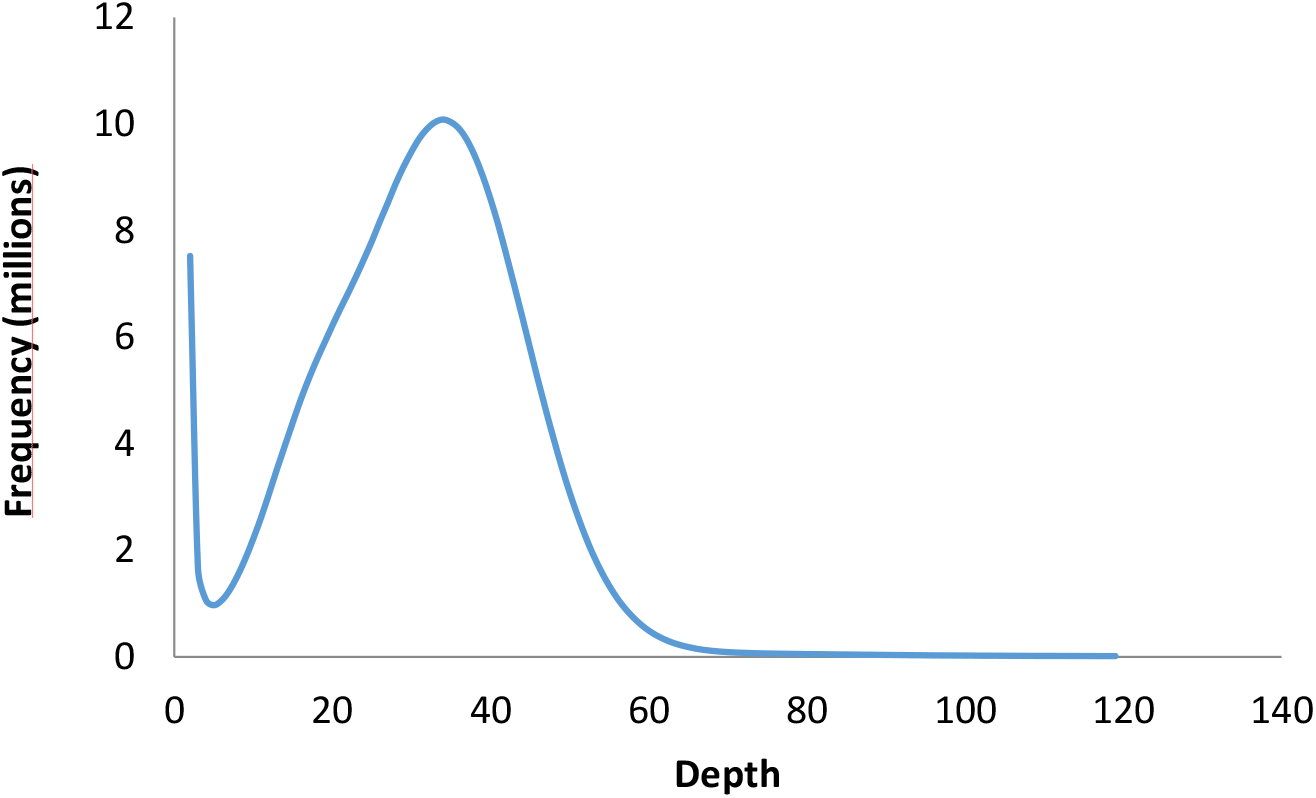
K-mer (K=31) distribution of Illumina genome sequencing reads of *R. maidis*. The total count of K-mers was 11,495,021,417, and the peak of K-mer depth was 34. The genome size of *R. maidis* was calculated by dividing the total K-mer count by the peak depth, which was approximately 338 Mb.

**Figure S2.**
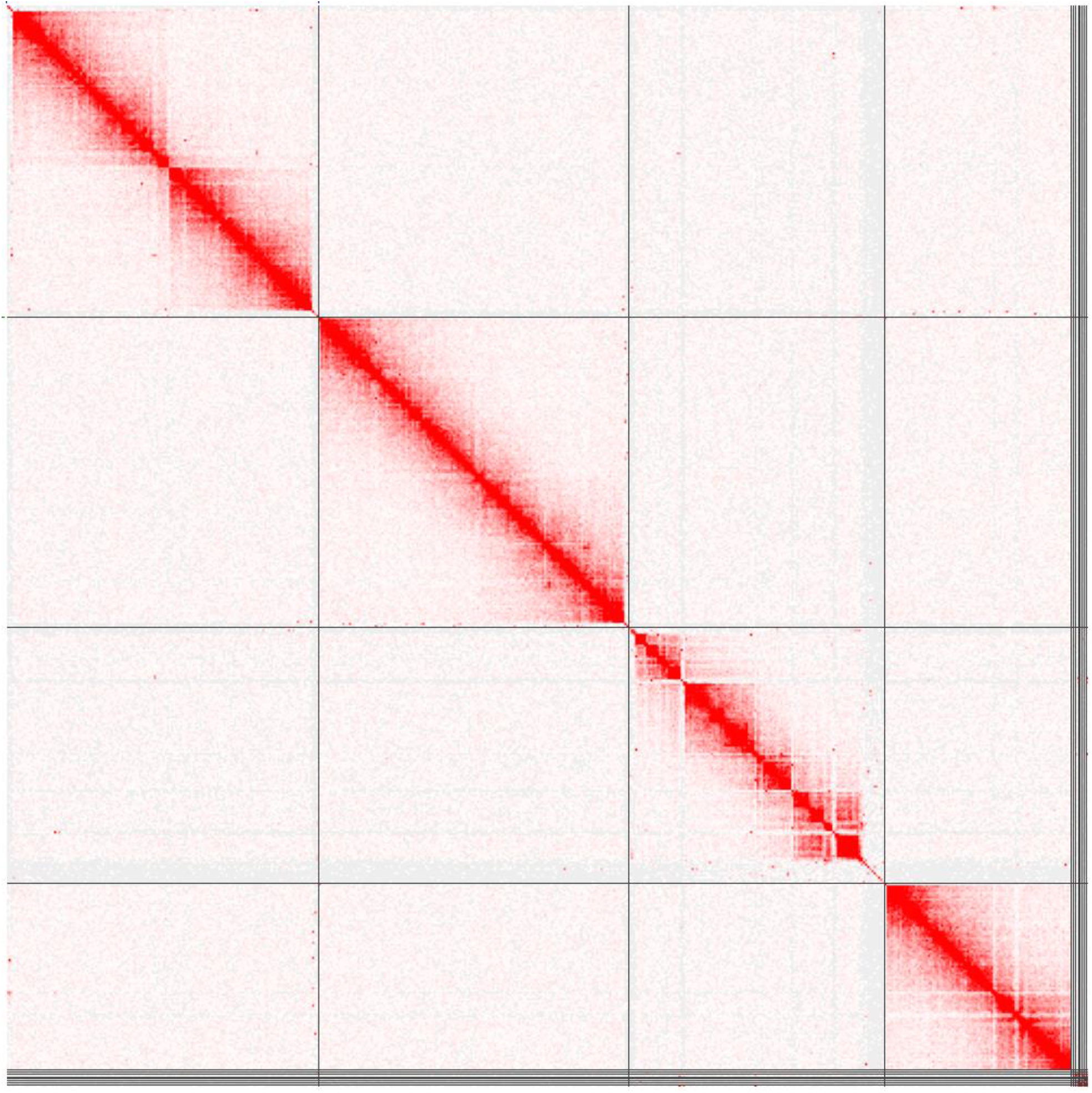
Hi-C contact map of the *R. maidis* genome

**Figure S3.**
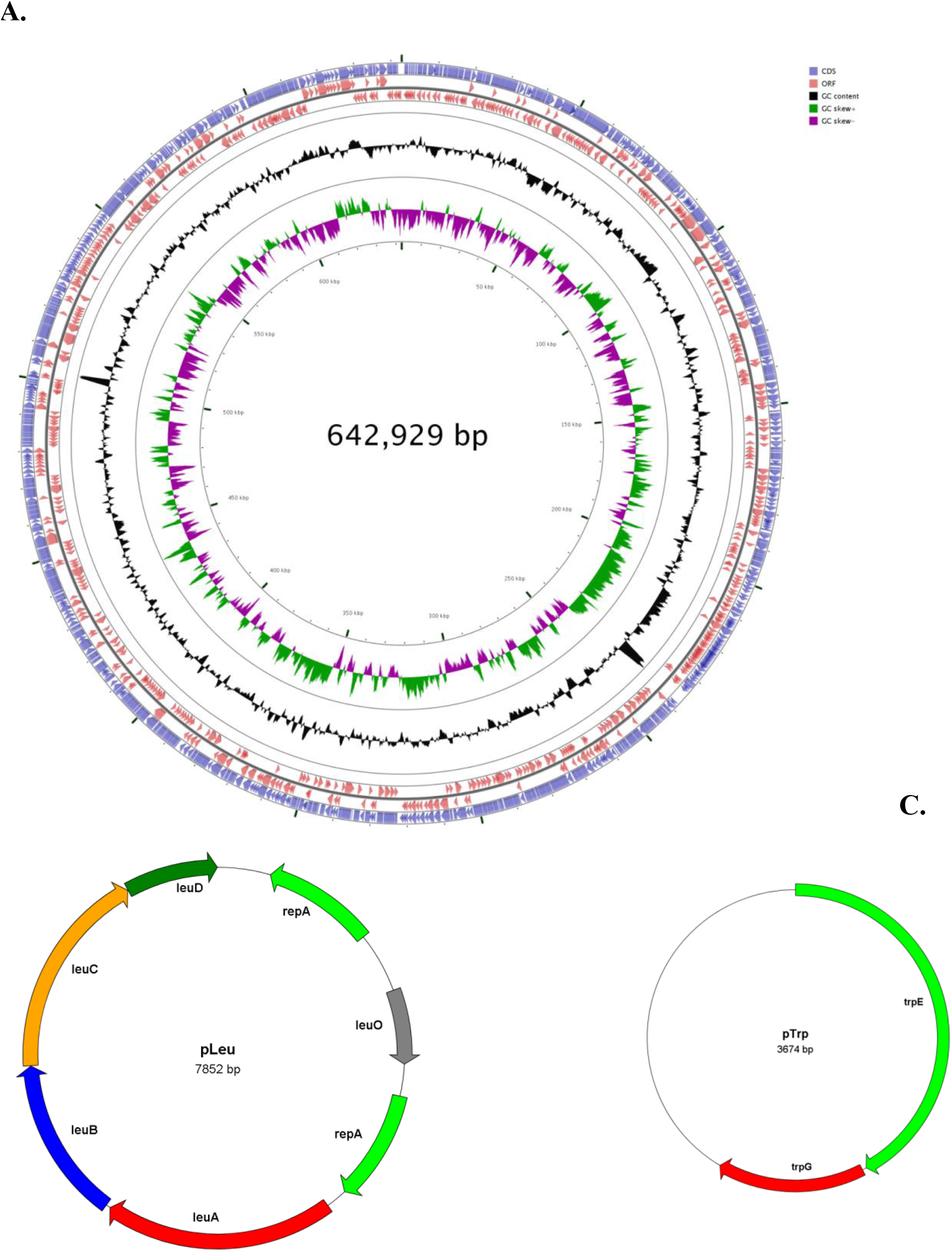
Circular view of the genome of the *Rhopalosiphum maidis* endosymbiont, *Buchnera aphidicola* (A) and its plasmids pLeu (B) and pTrp (C).

**Figure S4.**
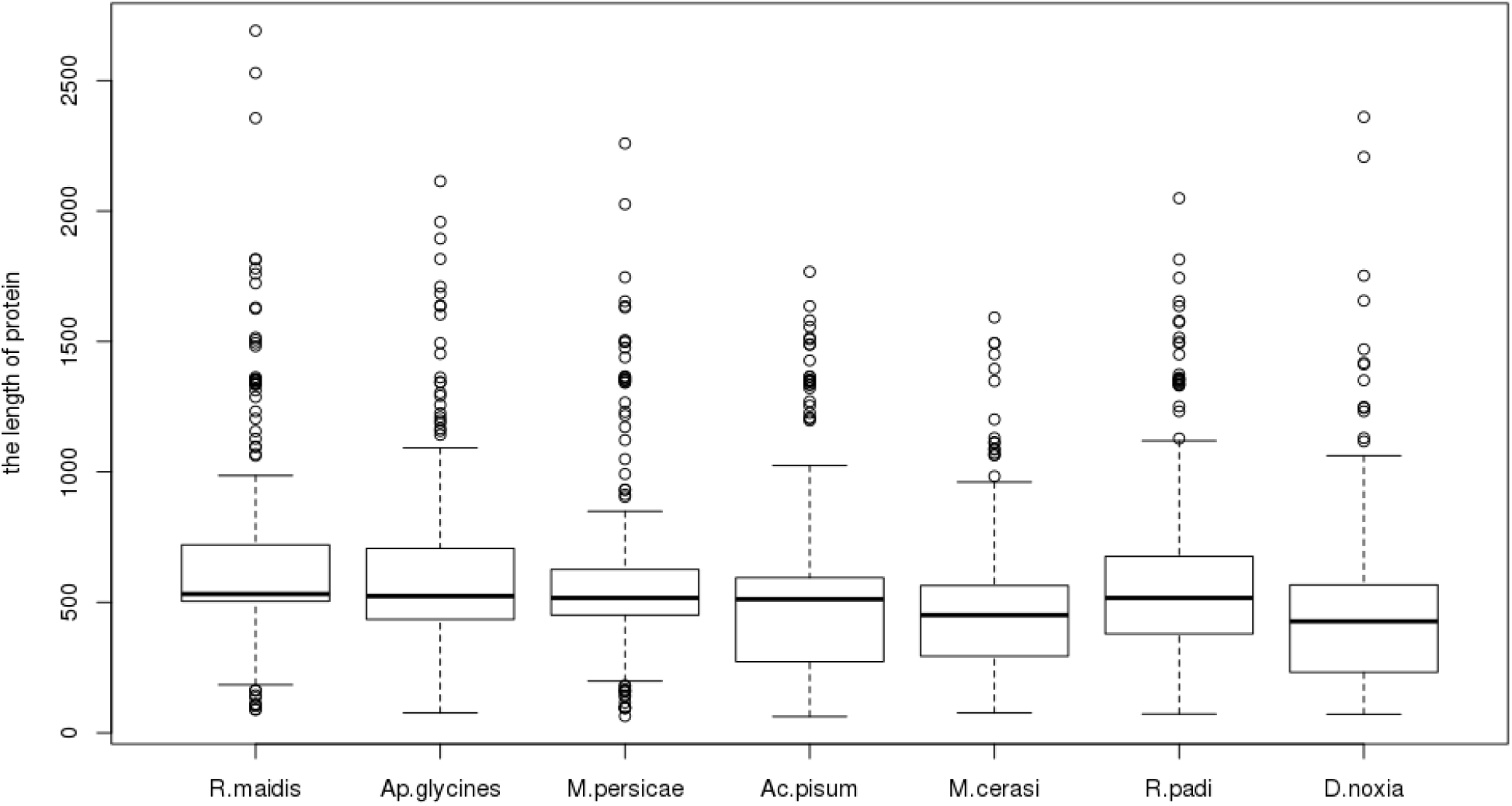
Length distribution of protein sequences of detoxification gene families in seven aphid species.

